# Multi-trait genome-wide association analyses leveraging alcohol use disorder findings identify novel loci for smoking behaviors in the Million Veteran Program

**DOI:** 10.1101/2022.10.18.512710

**Authors:** Youshu Cheng, Cecilia Dao, Hang Zhou, Boyang Li, Rachel L. Kember, Sylvanus Toikumo, Hongyu Zhao, Joel Gelernter, Henry R. Kranzler, Amy Justice, Ke Xu

## Abstract

Smoking behaviors and alcohol use disorder (AUD), moderately heritable traits, commonly co-occur in the general population. Single-trait genome-wide association studies (GWAS) have identified multiple loci for smoking and AUD. However, GWASs that have aimed to identify loci contributing to comorbid smoking and AUD have used small samples and thus have not been highly informative. Applying multi-trait analysis of GWASs (MTAG), we conducted a joint GWAS of smoking and AUD with data from the Million Veteran Program (N=318,694). By leveraging GWAS summary statistics for AUD, MTAG identified 21 genome-wide significant (GWS) loci associated with smoking initiation and 18 loci associated with smoking cessation compared to 16 and 8 loci, respectively, identified by single-trait GWAS. The novel loci for smoking behaviors identified by MTAG included those previously associated with psychiatric or substance use traits. Colocalization analysis identified 10 loci shared by AUD and smoking status traits, all of which achieved GWS in MTAG, including variants on *SIX3, NCAM1*, and near *DRD2*. Functional annotation of the MTAG variants highlighted biologically important regions on *ZBTB20, DRD2, PPP6C*, and *GCKR* that contribute to smoking behaviors. In contrast, MTAG of smoking behaviors and alcohol consumption (AC) did not enhance discovery compared with single-trait GWAS for smoking behaviors. We conclude that using MTAG to augment the power of GWAS enables the identification of novel genetic variants for commonly comorbid phenotypes, providing new insights into their pleiotropic effects on smoking behavior and AUD.

## Introduction

Smoking and alcohol use disorder (AUD) commonly co-occur in the general population. Compared to the use of a single substance, smoking comorbid with AUD has greater adverse health effects [1]. Smoking-related behaviors (e.g., smoking initiation, smoking cessation) and alcohol-related behaviors (e.g., alcohol consumption (AC) and AUD) have an estimated heritability of 40-50% [2-4]. The genetic correlations between smoking-related and alcohol-related behaviors are estimated to be about 40% [5, 6], suggesting that the pleiotropic effects of genetic variants contribute to their co-occurrence.

Genome-wide association studies (GWAS) with large sample sizes have made remarkable progress in identifying genetic loci for individual smoking-related and alcohol-related phenotypes. In a sample of over 1.2 million individuals, Liu et al. reported over 400 genome-wide significant (GWS) loci associated with multiple smoking-related and alcohol-related behaviors: 378 variants for smoking initiation, 24 variants for smoking cessation, and 99 variants for the number of alcoholic drinks consumed per week [7]. Quach et al. identified five loci for nicotine dependence in a meta-GWAS that included individuals with European ancestry and African ancestry [8]. In a sample of 209,915 European Americans (EA) from the Million Veteran Program (MVP), we reported 18 GWS loci for a smoking trajectory contrasting current versus never smoking (contrast I), which corresponds to smoking initiation, and five loci for another smoking trajectory contrasting current versus mixed smoking (contrast II), which is similar to smoking cessation [9]. Several dozen single nucleotide polymorphisms (SNPs) have been linked to alcohol misuse, AC, and AUD. For example, in another MVP study, we identified 13 loci for AC and 10 loci for AUD in an EA population [6]. In that study, as in others [10], AC and AUD were shown to have distinct genetic architectures.

Consistent with the phenotypic correlation between smoking-and alcohol-related behaviors, phenotypes related to these two different substances have moderate to strong genetic correlations [11, 12] and these correlations remain significant even after adjustment for environmental factors such as socioeconomic status [13]. However, the loci that contribute to the combined risks of smoking and drinking remain unclear, as standard GWAS considers traits in isolation rather than the combined influence of genetic variants on smoking and alcohol consumption or AUD. Thus, little is known regarding the pleiotropic effects of genetic variants on combined smoking-and alcohol-related phenotypes.

A recently developed method, multi-trait analysis of GWASs (MTAG), enables the joint analysis of genetically correlated traits to boost statistical power to detect variants for each trait [14]. MTAG takes summary statistics from single trait GWASs as input, generalizes inverse-variance-weighted meta-analysis to explore multiple traits, and calculates the trait-specific association for each variant. Moreover, MTAG accounts for overlap of samples among GWASs for different traits based on regression of linkage disequilibrium (LD) scores. Because of these features, MTAG has recently been applied to identify genetic variants for multiple, related phenotypes in psychiatric disorders and in substance use disorders. For example, Wu et al. utilized a sample size of approximately 60,000 EA individuals and identified genetic variants on seven genes commonly associated with four out of the following five psychiatric disorders: schizophrenia, bipolar disorder, autism spectrum disorder, attention-deficit hyperactivity disorder, and depression [15]. Recently, Deak et al applied MTAG to discover novel risk loci for opioid use disorder [16], and Xu et al. applied MTAG for four common substance use disorders and reported several novel loci for opioid use disorder, cannabis use disorder, alcohol and smoking behaviors [17]. These studies show that MTAG is a useful method for identifying loci associated with strongly correlated psychiatric disorders, including substance use behaviors.

In this study, we performed MTAG for two smoking-related behaviors (smoking initiation and smoking cessation) and two alcohol traits (AC, defined the same way as Alcohol Use Disorders Identification Test–Consumption (AUDIT-C), and AUD) in 318,694 EA individuals from the MVP database. Single-trait GWAS was performed for each of the four phenotypes, deriving summary statistics for MTAG. Multi-trait colocalization served to identify genetic risk loci shared by the smoking-and alcohol-related traits and to verify that MTAG augmented the power to identify the colocalized loci [18]. We also characterized MTAG performance by estimating the SNP-based heritability and heritability enrichment and by prioritizing causal genes. The analysis strategy is presented in **Figure 1**. Our results provide novel insight for the genetic contribution to the comorbidity of smoking-and alcohol-related behaviors.

**Figure 1:**
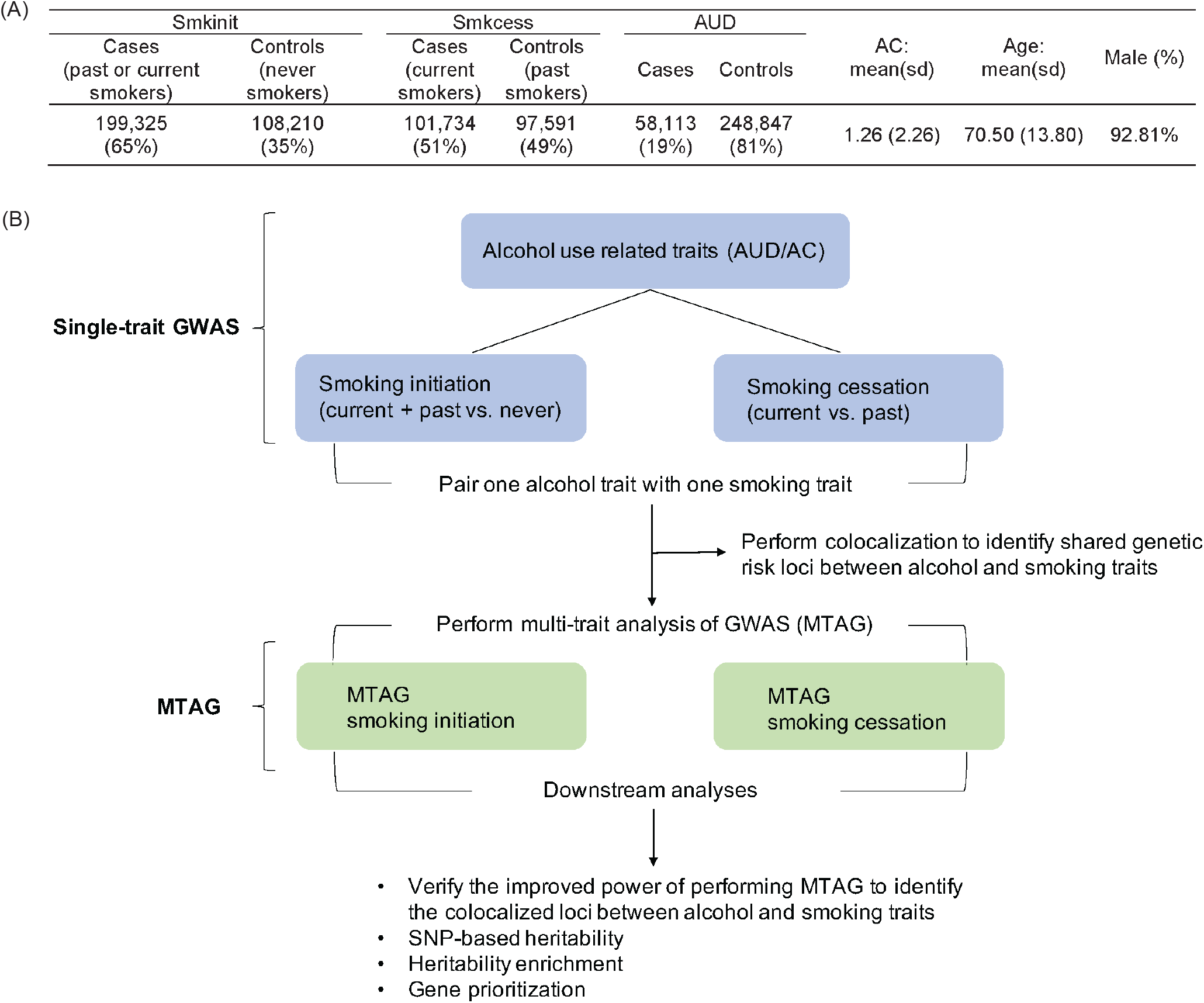
Phenotypes and analytic strategy. (A) The distribution of two smoking-related, two alcohol-related phenotypes, and demographic characteristics in the Million Veteran Program (MVP). (B) An overview of analyses performed on the single trait genome-wide association study (GWAS) and multi-trait GWAS (MTAG). All participants are from European American ancestry in the MVP (N = 318,694). Smkinit: smoking initiation; Smkcess: smoking cessation; AUD: alcohol use disorder; AC: alcohol consumption measured by Alcohol Use Disorders Identification Test-consumption (AUDIT-C).

## Methods

### Study samples and phenotypes

The MVP recruited veteran volunteers and collected data from them using questionnaires, access to their electronic medical records (EMRs), and genomic analysis of blood samples. The Institutional Review Board (IRB) of the Veterans Affairs Central Office and site-specific IRBs approved the MVP study. All relevant ethical regulations for work with human subjects were followed in the conduct of the study, and informed consent was obtained from all participants prior to data collection.

We used flashpca to perform principal component analysis (PCA) on all MVP samples and 2,504 samples from the 1,000 Genomes Project (1KG) to identify the genetic ancestry of subjects [19], which was unified with self-reported race/ethnicity using the HARE (Harmonizing Genetic Ancestry and Self-identified Race/Ethnicity) method to construct ancestral groups [20]. We removed samples with a high genotype missing rate (>10%), discordant sex, excessive heterozygosity (>3 SD), and up to second-degree relatives. A total of 318,694 EAs, 81,057 African Americans (AA), and 31,828 Hispanic Americans (HA) passed quality control filters. In the analyses reported herein, we focused on the MVP EA samples. Among the MVP EA samples, the mean age was 70.50 years with a standard deviation of 13.80. Most participants were male (92.81%).

Using the available EMR smoking observations, we identified 108,210 nonsmokers, 97,591 former smokers, and 101,734 current smokers. We defined smoking behaviors following Xu, Li, et al. [9]: individuals who ever smoked (former or current smokers) were contrasted with those who never smoked (nonsmokers) to study smoking initiation, while current smokers were contrasted with former smokers to explore smoking cessation.

For alcohol-related behaviors, we defined age-adjusted AUDIT-C (herein we call it AC) as described by Justice et al [21]. AUD cases were defined as individuals with ≥1 inpatient or ≥2 outpatient AUD codes according to the International Statistical Classification of Diseases and Related Health Problems, 9^th^ (ICD-9) or 10^th^ (ICD-10) revision; non-AUD (controls) were defined as the absence of any AUD code. The study sample comprised 58,113 individuals with AUD and 248,847 individuals without AUD.

### Genotyping and quality control

The MVP used an Affymetrix Axiom Biobank Array to genotype ∼ 723,000 markers. SNPs were validated for common diseases and phenotypes of specific interest to the VA population (e.g., psychiatric traits) [22]. Minimac4 and the 1000 Genomes Project 3 reference panel were used to conduct genotype imputation [23, 24]. During the quality control step, we filtered out variants that were rare (minor allele frequency < 0.01), had a missing rate > 5%, an imputation r^2^ < 0.8, or that deviated significantly from Hardy– Weinberg equilibrium (p < 1E−6). This yielded a total of 4.14 million variants.

### Single trait GWAS for smoking and alcohol traits

For the smoking phenotypes (smoking initiation and smoking cessation) and AUD, logistic regression was applied to estimate marginal effects of each single genetic variant on the phenotype, while for AC, linear regression was used. PLINK (v1.9) was employed to perform logistic and linear regression analyses [25], and age, sex, and the top 10 genotype principal components (PCs) calculated by flashpca were included in each model as covariates.

### MTAG analysis for smoking-related behaviors

We used MTAG to jointly analyze summary statistics of one alcohol-related GWAS with one smoking-related GWAS [14], yielding a total of 4 combinations: AUD with smoking initiation, AUD with smoking cessation, AC with smoking initiation, AC with smoking cessation. The MTAG results for smoking behaviors were our focus.

Lead SNPs and risk loci were defined in the same way for the single-trait GWAS and MTAG summary statistics: independent SNPs (LD, r^2^ < 0.1) with the most significant p-values were identified as lead SNPs, while the region containing all GWS variants (p < 5E−8) that were in LD (r^2^ > 0.6) with the lead SNP was defined as a risk locus. We merged loci within 250 kb [26, 27]. ANNOVAR was then employed to map lead SNPs to their nearest genes [28], and loci whose lead SNPs were mapped to the same gene were further merged into a single risk locus [9].

### Colocalization between AUD and smoking-related behaviors

To identify genetic risk factors shared by AUD and smoking-related behaviors, we applied HyPrColoc (Hypothesis Prioritization for multi-trait Colocalization) [18] in multiple genomic regions using summary statistics from single-trait GWAS. HyPrColoc reports (1) the posterior probability that alcohol and smoking behaviors are colocalized in a specific region, (2) the causal variant in this colocalized region and the proportion of the posterior probability of colocalization explained by this variant. The identified shared risk variants facilitate validation of whether MTAG improves the power to identify colocalized loci between alcohol and smoking traits. We first used LDetect to partition the genome into 2258 independent regions (each, on average, approximately 1.6 cM in length) [29, 30], with LD estimated from the 1000 Genomes Project phase III samples of European ancestry [31]. We paired AUD with each of the smoking behaviors and performed colocalization analysis to identify shared genetic risk factors. For each pair, we reported the regions whose posterior probability of colocalization was greater than 0.75, as suggested by the authors of HyPrColoc [18]. LocusZoom (v1.3) was applied to visualize the change in regional associations after performing MTAG [32].

### Downstream analysis of the results from single-trait GWAS and MTAG

We estimated heritability and heritability enrichment for the 2 smoking-related single-trait GWASs and 2 smoking-related MTAGs. LD score regression (v1.0.0) was performed to estimate the narrow-sense heritability due to additive genetic effects [33]. To identify tissues most relevant to smoking-related phenotypes, we performed heritability enrichment analyses using 66 functional annotations from GenoSkyline-Plus (v1.0.0), which included tissues and cell lines from the blood, brain, lung, vascular system, heart, thymus, spleen, muscles, gastrointestinal tract, pancreas, liver, fat, bone/connective tissue, skin, breast, and ovary [34]. To adjust for multiple comparisons, we applied Bonferroni correction to the 66 enrichment tests for 2 smoking-related single-trait GWASs and 2 smoking-related MTAGs (p < 0.05/66/4 = 1.89E−4).

Functional gene mapping was performed for 2 smoking-related MTAGs. We used the FUMA tool (v1.3.6) to conduct eQTL and chromatin interaction mapping [35]. We restricted eQTL mapping to 13 genotype-tissue expression (GTEx) v8 brain tissues and performed chromatin interaction mapping with the built-in adult cortex Hi–C data and enhancer/promoter annotations in 12 brain tissues from the Roadmap epigenomes. By default, we used false-discovery rate (FDR) < 0.05 for significant SNP-gene pairs in the eQTL mapping and FDR < 1E−6 for significant chromatin interactions, as suggested by Schmitt et al [36].

## Results

### Single-trait GWAS for smoking initiation, cessation, and alcohol behaviors

*Single-trait GWAS for smoking initiation and cessation*: We previously reported 12 GWS loci associated with smoking initiation and 8 loci associated with smoking cessation among 209,915 EA individuals in the MVP database [9]. In this study, where we utilized a larger sample from the MVP (N=318,694), we identified 16 loci for smoking initiation (**Supplementary Figure 1A**) (**Supplementary Table 1**), five of which were previously reported to be linked to smoking initiation (*TEX41, ZBTB20, EPHX2, NCAM1*, and *SPATS2*), including two identical lead SNPs, rs6438208 on *ZBTB20* and rs78875955 on *EPHX2*. In addition, we identified novel loci for smoking initiation, which included several RNA coding genes: *Y_RNA, LINC01360, and LINC01833*. The intronic SNP rs4687552 on *ITIH3* was also a novel locus associated with smoking initiation in this sample.

We found eight GWS loci associated with smoking cessation, including two that we previously reported: rs6011779 on *CHRNA4* and rs17602038 on *DRD2* (**Supplementary Figure 1B**) (**Supplementary Table 1**). We also previously reported a GWS association of rs112270518 near *DBH* with smoking trajectory II (current versus mixed smoking) [9], which corresponds to the smoking cessation trait. Here, rs3025360, near *DBH*, was a GWS locus associated with smoking cessation. Rs12341778 on *MAPKAP1* and rs11881918 near *CYP2A6*, with the mapped genes previously linked to mood disorder [37] and nicotine metabolite ratio [38] respectively, also showed GWS associations with smoking cessation. The other two novel GWS loci associated with smoking cessation in the present study were 3:49638084:A:AAAATT on *BSN* and rs650599 on *SCAI*.

*Single-trait GWAS for alcohol phenotypes*: Compared with our previous GWASs on AUD and AC in the MVP cohort [6, 39], here with a larger MVP sample size, we identified eight loci that overlapped with previously reported loci for AUD (i.e., *GCKR, SIX3, ARHGAP15, ADH1B, SLC39A8, CNTLN, DRD2*, and *FTO*) as well as six novel loci for AUD (*LNC01360, FANCL, LOC646736, PLCL2, KLB*, and *MTCH2*). Regarding AC, there were 27 GWS loci, six of which overlapped with loci associated with AUD, including five that we previously reported. Rs13130101, near *KLB*, was a GWS locus associated with AUD, and rs13146907, also near *KLB*, was a GWS locus associated with AC (**Supplementary Table 1** and **Supplementary Figure 1C, 1D**). *KLB* is a coreceptor for the hormone FGF21 and was previously linked to alcohol intake in a European ancestry population [40]. Altogether, we identified more loci for each smoking-and alcohol-related trait in the present study with a larger sample size drawn from the MVP database.

*Genetic correlations between smoking-and alcohol-related phenotypes*: The genetic correlation between smoking behaviors and AUD ranged from 0.59 to 0.62, substantially higher than the correlation between smoking behaviors and AC (ranging from 0.08 to 0.12) (**Supplementary Figure 2**) (**Supplementary Table 1**). This pattern indicates that the shared genetic risk between smoking-related phenotypes and AUD is much greater than that between smoking-related phenotypes and AC.

**Figure 2:**
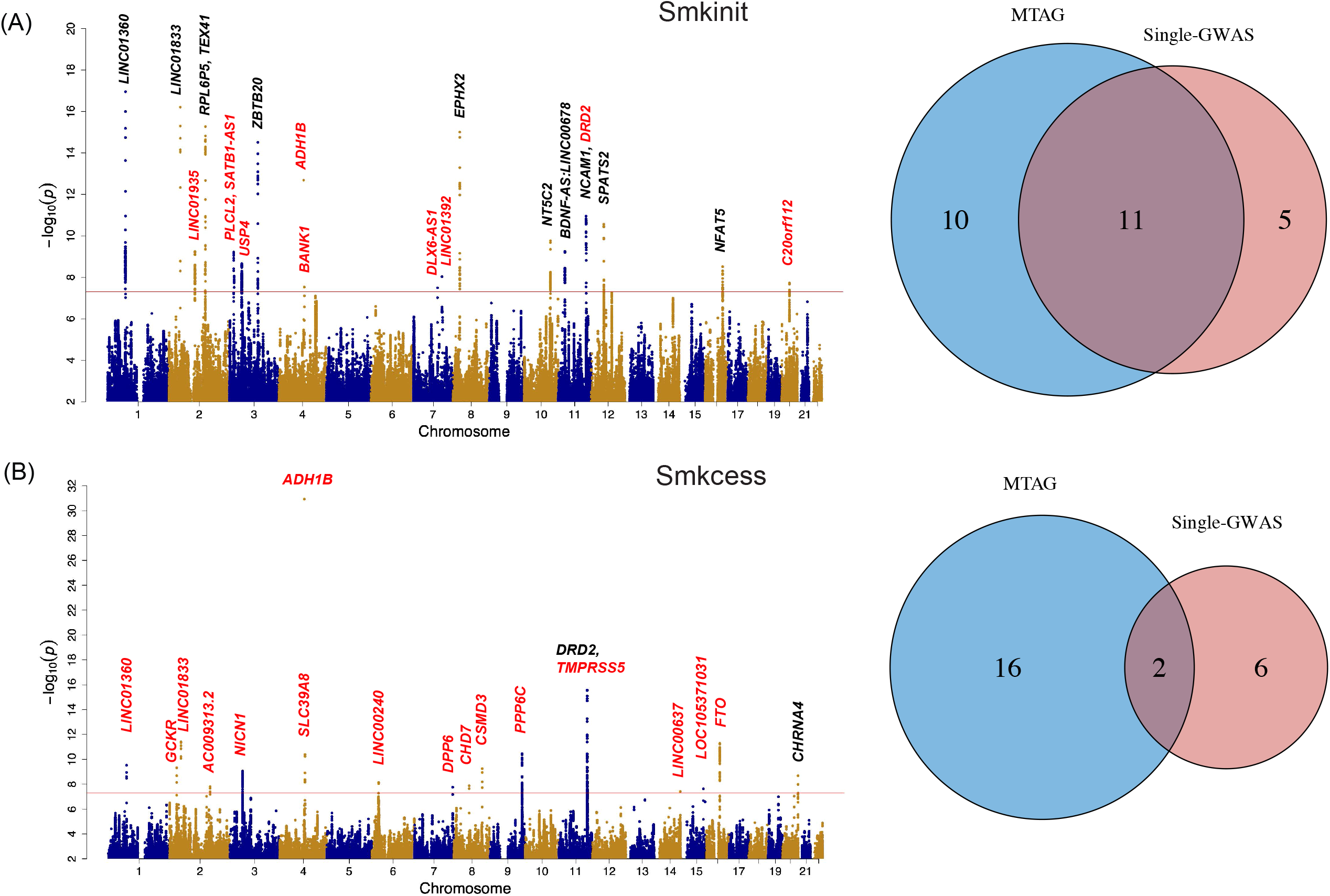
Multi-trait analysis of GWASs (MTAG) on two smoking phenotypes with alcohol use disorder (AUD). Manhattan plot of MTAG and Venn plot of the number of genome-wide significant (GWS) loci identified by single-trait GWAS and MTAG for (A) smoking initiation and (B) smoking cessation. The nearest genes to GWS loci are shown. New MTAG-identified loci are shown in red. Smkinit: smoking initiation; Smkcess: smoking cessation; AUD: alcohol use disorder.

### MTAG analysis for smoking-related behaviors

We conducted a joint-GWAS for two smoking-related phenotypes (smoking initiation, smoking cessation) and two alcohol-related phenotypes (AUD and AC) using MTAG. Each smoking phenotype was paired with AUD or AC.

Leveraging the summary statistics from single-trait GWAS for smoking initiation and AUD, MTAG identified 21 GWS loci for smoking initiation; five more than the number of loci identified through single-trait GWAS for smoking initiation. Among the 21 loci, 11 were identified by the single-trait GWAS for smoking initiation, while 10 were novel loci from the MTAG results (**Figure 2A**). The novel loci identified by MTAG included the well-known loci for alcohol phenotypes, rs1229984 on *ADH1B* and rs6589386 on *DRD2* (**Supplementary Table 2**). Another novel locus for smoking initiation has been linked with smoking-related phenotypes in previous studies: rs6778080 on *USP4* was linked to lifetime smoking index and depression [41, 42].

Regarding smoking cessation, MTAG identified 18 GWS loci, 10 more than the single-trait GWAS for smoking cessation. Two of the 18 loci, rs6011779 on *CHRNA4* and rs17602038 on *DRD2*, were identified by both the single-trait GWAS and MTAG for smoking cessation. Sixteen of these loci attained GWS only in MTAG (**Figure 2B**). As with the MTAG for smoking initiation, we found that multiple alcohol-related loci also attained GWS for smoking cessation. These loci included rs1229984 on *ADH1B* and rs62048402 on *FTO* (**Supplementary Table 2**). Another highly pleiotropic locus, rs13135092 on *SLC39A8*, previously associated with high-density lipoprotein cholesterol (HDL) in current drinkers [43] and schizophrenia [44], was a GWS locus associated with smoking cessation. Additionally, rs1260326 on *GCKR*, which has been linked to multiple metabolic traits [45], attained GWS for smoking cessation.

Of note, compared with the single-trait GWAS results (**Supplementary Figure 1A, 1B**), the MTAG of AC and smoking phenotypes did not yield new loci for smoking phenotypes (**Supplementary Figure 3**). The subsequent analyses included only the results from the MTAG of each smoking trait and AUD. Together, these findings show that by leveraging the strongly correlated AUD trait, MTAG was able to identify more loci for smoking phenotypes than single-trait GWAS.

**Figure 3:**
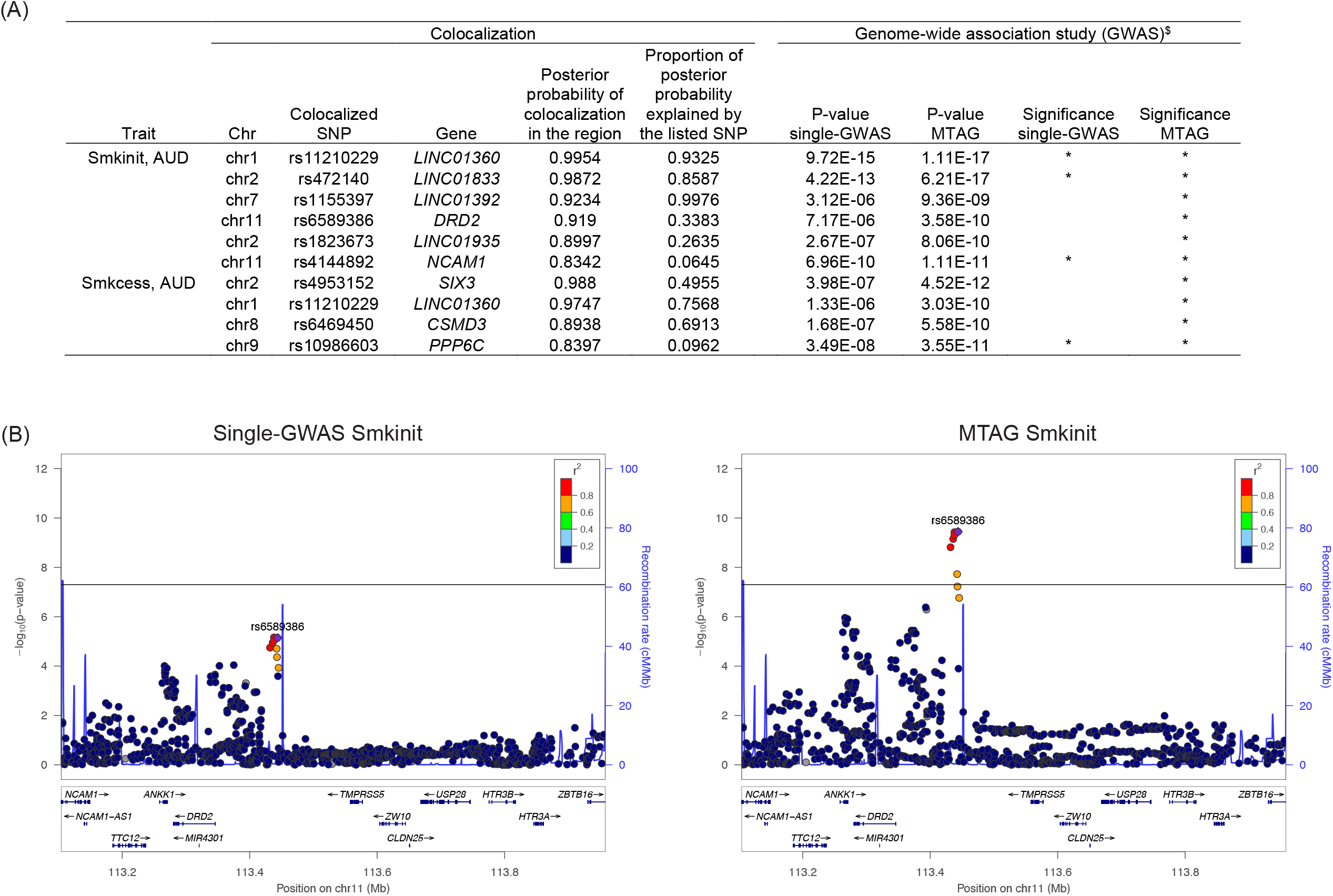
Multi-trait colocalization analysis of two smoking traits and alcohol use disorder (AUD). (A) Hypothesis prioritization for multi-trait colocalization (HyPrColoc) identified six regions shared by AUD and smoking initiation and four regions shared by AUD and smoking cessation. We report regions whose posterior probability of colocalization was greater than 0.75. ^$^: p values for the colocalized SNP in the single-trait GWAS and in MTAG for the corresponding smoking trait. (B) LocusZoom plots for the association of rs6589386 with smoking initiation. The genetic variant rs6589386 mapped near *DRD2* was identified as a colocalized SNP between AUD and smoking initiation. Smkinit: smoking initiation; Smkcess: smoking cessation; AUD: alcohol use disorder.

### Colocalization between AUD and smoking-related behaviors

To identify genetic risk factors shared by AUD and smoking-related behaviors, we performed hypothesis prioritization for multi-trait colocalization (HyPrColoc) by pairing single-trait GWAS for each of the two smoking traits with single-trait GWAS for AUD. For each pair, we reported the regions whose posterior probability of colocalization was greater than 0.75 [18]. We identified a total of 10 colocalized regions, including six regions shared by AUD and smoking initiation and four shared by AUD and smoking cessation (**Figure 3A**). Of note, among the 10 regions shared by AUD and smoking traits, MTAG identified all as attaining GWS, while single-trait GWAS identified only 4 out of 10 as GWS loci. Thus, the greater power of MTAG is most obvious for loci shared by AUD and smoking-related behaviors. For example, one colocalized SNP for AUD and smoking initiation, rs6589386 on *DRD2*, was marginally significant in the single-trait GWAS (p-value = 7.17E-06) but attained GWS in MTAG (p-value = 3.58E-10) (**Figure 3B**).

Another noteworthy colocalized SNP shared by AUD and two smoking traits was rs11210229 on *LINC01360* (**Supplementary Figure 4**). For smoking initiation, MTAG resulted in a moderate increase in the significance of the association with rs11210229 and other variants in LD. For smoking cessation, the increase in significance was greater: rs11210229 did not attain significance in the single-trait GWAS (p-value = 1.33E-06) but was the lead GWS SNP in MTAG (p-value = 3.03E-10).

**Figure 4:**
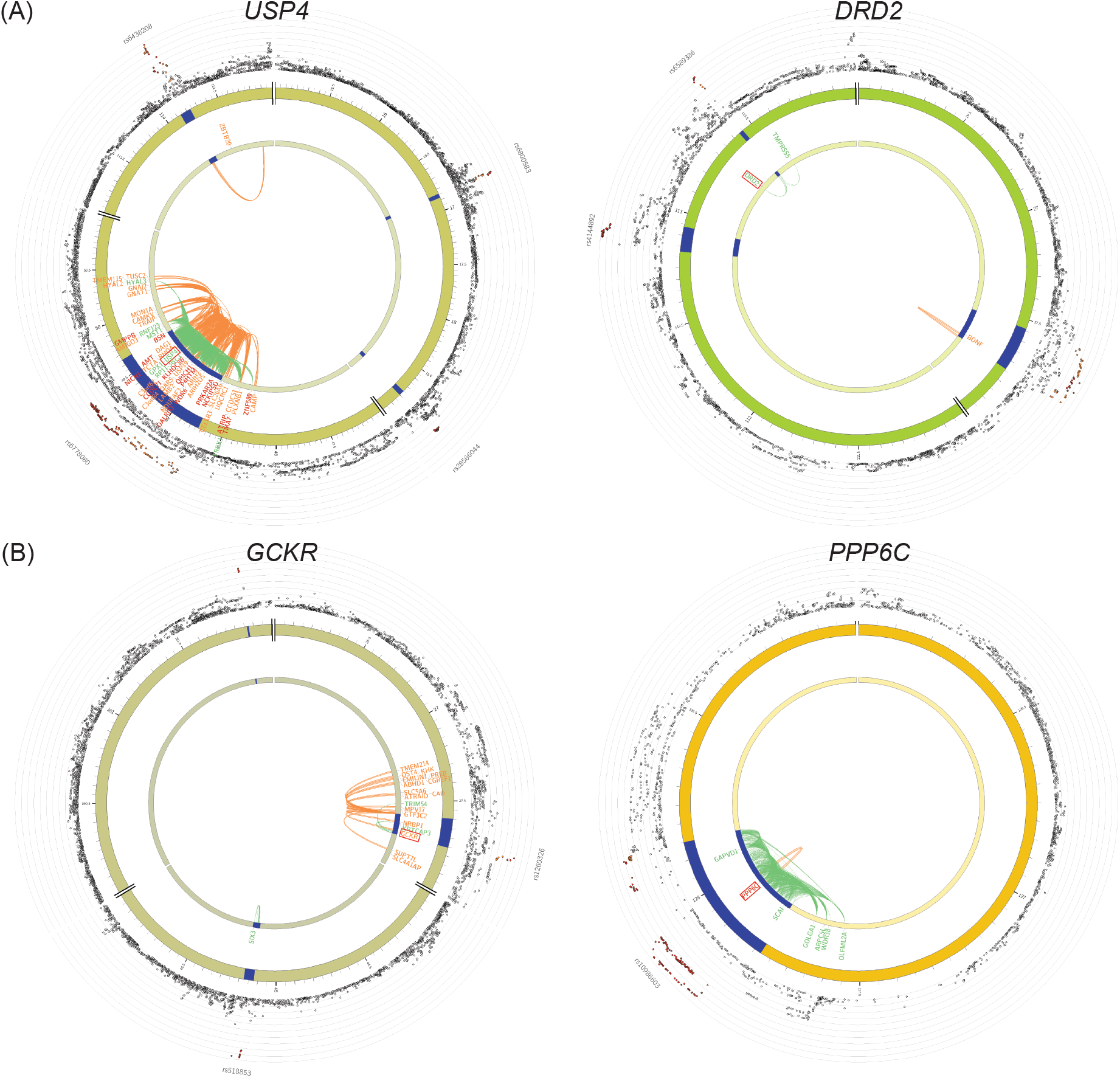
Gene prioritization for smoking traits using the MTAG. Functional mapping and annotation (FUMA) gene prioritization were performed for (A) smoking initiation and (B) smoking cessation. The outer layer shows chromosomal Manhattan plots. The GWS locus is indicated in blue. Genes mapped by chromatin interactions or eQTL are shown in orange or green, respectively. Genes mapped by both chromatin interactions and eQTL are shown in red.

### Estimated heritability and enrichment for smoking-related behaviors

The estimated heritability from MTAG was 1-2% greater for each smoking trait than the heritability from single-trait GWAS (**Supplementary Table 3**). For the two smoking traits, single-trait GWAS did not detect significant heritability enrichment, while MTAG identified significant heritability enrichment for smoking initiation in the anterior caudate (enrichment = 4.08, Wald test p = 1.86E-05) and the dorsolateral prefrontal cortex (enrichment = 5.41, Wald test p = 4.40E-05) as well as significant heritability enrichment for smoking cessation in the anterior caudate (enrichment = 5.64, Wald test p = 2.72E-06) and the colonic mucosa (enrichment = 7.69, Wald test p = 4.61E-05) (**Supplementary Table 3**).

### Prioritizing genetic regions for smoking phenotypes based on MTAG

By integrating MTAG variants with functional genomic features in brain tissues, we identified biologically important regions/genes for smoking behaviors. We performed expression quantitative trait loci (eQTL) and chromatin interaction mapping for the MTAG summary statistics of two smoking-related behaviors with the functional mapping and annotation (FUMA) tool [35]. For smoking initiation, we identified 41 genes mapped by eQTL and 85 genes mapped by chromatin interaction (**Supplementary Table 4**). Among those regions, seven overlapped with MTAG-identified loci, including two newly identified loci, rs6589386 on *DRD2* and rs6778080 on *USP4* (**Figure 4A**); this finding indicates the causal role of those genes. For smoking cessation, we identified 46 significant genomic regions by eQTL mapping and 67 regions by chromatin interaction mapping. Importantly, six of those regions overlapped with MTAG-identified loci, including five newly identified loci (**Supplementary Table 4**). For example, the MTAG-identified SNP rs10986603 on *PPP6C* colocalized with *PPP6C* eQTL in the anterior cingulate cortex and chromatin interaction in the cortex (**Figure 4B**). *PPP6C* was recently associated with opioid addiction in EA individuals according to gene-based and eQTL analyses [46]. This finding was replicated in a GWAS on opioid use disorder conducted among MVP participants [47].

## Discussion

By leveraging the genetic architecture of AUD, we identified new loci for smoking behaviors that were not identified using a single-trait GWAS approach. MTAG revealed genetic variants that affect both smoking behaviors and AUD. Convergent evidence from MTAG, colocalization, and functional annotation analyses highlighted several AUD-associated genes that contribute to smoking behaviors. Importantly, the newly identified loci for smoking were colocalized with eQTL and chromatin interaction in brain regions previously shown to be relevant for addictive behavior. These findings underscore prior findings that MTAG is a powerful approach for identifying genetic variants for complex traits, as it is particularly important for revealing pleiotropic effects of significant variants that contribute to highly comorbid disorders. Although smoking and alcohol use are closely associated both epidemiologically and clinically, our study revealed GWS variants contributing to the cooccurrence of these traits in a large EA population, thereby providing insight into their pleiotropic effects on the comorbid phenotypes.

Using MTAG, we replicated previous findings that linked genomic regions to smoking initiation and smoking cessation in the Genetic Sequencing Consortium of Alcohol and Nicotine Use (GSCAN) study of 1.2 million individuals [7]. Among the 21 MTAG-identified variants for smoking initiation, nine had the same nearest genes as those in the GSCAN study, including *SPATS2L, PLCL2, BDNF-AS*, and *NCAM1*. For smoking cessation, five of 18 MTAG-loci overlapped with the GSCAN study, including three long intergenic nonprotein coding RNA genes (i.e., *LINC01360, LINC01833, and LINC00637*). However, AUD-associated genes such as *ADH1B* and *FTO* only attained GWS for smoking initiation or smoking cessation in the present study.

This study provides novel genomic findings that link well-established genetic loci for AUD to smoking behavior. These findings augment well-established phenotypic associations showing both high rates of smoking among individuals with AUD [48] and greater difficulty in stopping smoking among individuals with AUD [49]. A functional variant, rs1229984 on *ADH1B*, has long been recognized as a risk locus for AC and alcohol-related diseases across populations with different ancestry. Recently, this locus, combined with another functional locus on *ALDH2*, was shown to be predictive of smoking initiation in a Japanese population [50]. We found that rs1229984 was strongly associated with two smoking behaviors in the context of AUD, suggesting that rs1229984 influences smoking behavior for individuals with problematic alcohol use.

In addition, rs9937709, near *FTO*, was significantly associated with smoking cessation. *FTO* has been linked to obesity [51], AC, and AUD [6]. We previously reported that rs62033408, a lead SNP on *FTO*, was associated with AC and that rs1421085, near *FTO*, was associated with AUD. The MTAG results indicated that rs9937709, near *FTO* and 19.9 kb from rs1421085, was associated with smoking behaviors. Another region near *DRD2* was important for alcohol and smoking behaviors: we previously reported significant associations of rs61902812 with AUD and of rs3133388 with smoking trajectory contrast I (current versus never smoking, corresponding to smoking initiation). In this study, MTAG identified multiple novel loci near *DRD2* for different smoking behaviors, including rs6589386 for smoking initiation and rs17602038 for smoking cessation, suggesting that this region is biologically important for understanding the mechanism underlying how variation in *DRD2* leads to addictive behaviors.

In contrast to the highly informative MTAG for AUD and smoking behaviors, the MTAG for AC and smoking behaviors did not show enhanced effects relative to the single-trait GWAS. Although the reasons for these differences are unclear, we hypothesize that the different genetic architectures of AC and AUD partially explain these findings. As we previously reported, AC and AUD have distinct profiles of genetic correlations [6, 10]. AC is negatively genetically correlated with some medical diseases, such as coronary artery disease and Type 2 diabetes, while AUD is positively genetically correlated with psychiatric diseases, including addictive disorders. Consistent with other studies, we found that the genetic correlations between AUD and the smoking behaviors were greater than those between AC and the smoking behaviors. In the MTAG results presented here, the genetic correlations between AUD and the different smoking behaviors were approximately 0.6, while those between AC and the smoking behaviors were approximately 0.1. This pattern is consistent with the original report [14], which emphasized that MTAG is most useful for analyzing phenotypes with strong genetic correlations. Another explanation is that the AC measurements may have been inaccurate. The AUDIT-C component of the MVP study is a self-reported measure of alcohol intake over the past 12 months. As we previously reported, individuals who were lifetime abstainers or former drinkers could have a confounding effect on gene associations with AC [52]. After removing former alcohol drinkers, single-trait GWAS identified more loci for AC in a sample from the UK Biobank. Future studies using longitudinal assessment of AC are warranted to deepen our understanding of the genetic architecture of smoking behaviors in the context of AUD and AC.

We acknowledge several limitations in the study. A lack of ancestral diversity limits the findings to EA individuals, due to lack of availability of GWASs on non-EA individuals that are large enough to provide adequate statistical power. As larger samples of other ancestral groups become available, we plan to examine whether the findings reported here are replicable in other populations. Second, MTAG is established on the assumption that all SNPs share the same variance–covariance matrix of effect sizes across multiple traits [14]. This assumption may not be applicable to AUD and smoking behavior. Authors of MTAG also stressed one potential problem for SNPs that are true null for one trait but non-null for another trait. For such SNPs, MTAG could have false positives in the first trait [14]. This statement is consistent with a recent study on opioid use disorder, which showed that increased detection of MTAG might come with the loss of specificity [16]. In our analyses, some of the newly identified loci by MTAG were verified to be shared between AUD and smoking behaviors by colocalization, such as rs6589386 on *DRD2*. However, there also existed MTAG-identified loci that were not significant in the colocalization analysis, such as rs1229984 on *ADH1B*, suggesting that some loci identified by MTAG may be biased towards AUD. The MTAG-identified rs1229984 on *ADH1B* for smoking traits need to be interpreted with cautiousness. Further investigations are needed to verify whether this locus revealed by MTAG is truly shared across different phenotypes. Third, we were unable to differentiate lifetime abstainers from former alcohol users who quit drinking (possibly due to alcohol-related problems) in the MVP samples. Future studies that use biomarkers instead of self-reported data to quantify AC could benefit from greater statistical power to detect risk loci for smoking behaviors. Finally, we did not examine the functional effects of the pleiotropic loci for smoking and AUD; such studies are needed to understand the mechanisms underlying the findings we report.

In summary, we identified multiple genetic loci significantly associated with the cooccurrence of smoking behavior and AUD in an EA population. The findings highlight several biologically relevant regions for further study that could elucidate mechanisms shared by smoking behavior and AUD and thus provide opportunities for novel interventions that target these mechanisms.

## Acknowledgement

This research is based on data from the Million Veteran Program, Office of Research and Development, Veterans Health Administration, and was supported by award I01 BX004820 (MVP004) and the Veterans Integrated Service Network 4 Mental Illness Research, Education and Clinical Center at the Crescenz VAMC. The study was also supported by National Institute on Drug Abuse grants R01 DA038632, R01 DA047063, and R01 DA047820. This publication does not represent the views of the Department of Veterans Affairs or the United States Government. We thank all participants in the MVP who allowed access to their electronic health records and provided blood samples for genomic analyses. We thank all authors and consortia whose data were used here for making their summary statistics publicly available to the research community.

## Author contributions

YC, CD and KX contributed to the analytic plan. CD performed the single-trait GWAS analyses. YC performed MTAG, colocalization and downstream analyses. The Million Veterans Program provided data and computational platform. HZhou, BL, RLK, ST, HZhao, JG, HRK, AJ, KX contributed to data interpretation. YC and KX prepared the first version of the manuscript. CD, HZhou, BL, RLK, ST, HZhao, JG, HRK, AJ contributed to manuscript preparation and all authors approved the paper.

## Additional information

Supplementary information is available at MP’s website

## References

1. Adams S. Psychopharmacology of Tobacco and Alcohol Comorbidity: a Review of Current Evidence. Curr Addict Rep. 2017;4(1):25–34.

2. Verhulst B, Neale MC, Kendler KS. The heritability of alcohol use disorders: a metaanalysis of twin and adoption studies. Psychol Med. 2015;45(5):1061–72.

3. Vink JM, Willemsen G, Boomsma DI. Heritability of smoking initiation and nicotine dependence. Behav Genet. 2005;35(4):397–406.

4. Carmelli D, Swan GE, Robinette D, Fabsitz R. Genetic influence on smoking--a study of male twins. N Engl J Med. 1992;327(12):829–33.

5. Clarke TK, Adams MJ, Davies G, Howard DM, Hall LS, Padmanabhan S, et al. Genomewide association study of alcohol consumption and genetic overlap with other health-related traits in UK Biobank (N=1121117). Mol Psychiatry. 2017;22(10):1376–84.

6. Kranzler HR, Zhou H, Kember RL, Vickers Smith R, Justice AC, Damrauer S, et al. Genomewide association study of alcohol consumption and use disorder in 274,424 individuals from multiple populations. Nature Communications. 2019;10(1):1499.

7. Liu M, Jiang Y, Wedow R, Li Y, Brazel DM, Chen F, et al. Association studies of up to 1.2 million individuals yield new insights into the genetic etiology of tobacco and alcohol use. Nat Genet. 2019;51(2):237–44.

8. Quach BC, Bray MJ, Gaddis NC, Liu M, Palviainen T, Minica CC, et al. Expanding the genetic architecture of nicotine dependence and its shared genetics with multiple traits. Nat Commun. 2020;11(1):5562.

9. Xu K, Li B, McGinnis KA, Vickers-Smith R, Dao C, Sun N, et al. Genome-wide association study of smoking trajectory and meta-analysis of smoking status in 842,000 individuals. Nature Communications. 2020;11(1):5302.

10. Sanchez-Roige S, Palmer AA, Clarke TK. Recent Efforts to Dissect the Genetic Basis of Alcohol Use and Abuse. Biol Psychiatry. 2020;87(7):609–18.

11. Madden PA, Bucholz KK, Martin NG, Heath AC. Smoking and the genetic contribution to alcohol-dependence risk. Alcohol Res Health. 2000;24(4):209–14.

12. Zhang T, Gao W, Cao W, Zhan S, Lv J, Pang Z, et al. The Genetic Correlation Between Cigarette Smoking and Alcohol Drinking Among Chinese Adult Male Twins: An Ordinal Bivariate Genetic Analysis. Twin Research and Human Genetics. 2012;15(4):483–90.

13. Marees AT, Smit DJA, Abdellaoui A, Nivard MG, van den Brink W, Denys D, et al. Genetic correlates of socio-economic status influence the pattern of shared heritability across mental health traits. Nat Hum Behav. 2021;5(8):1065–73.

14. Turley P, Walters RK, Maghzian O, Okbay A, Lee JJ, Fontana MA, et al. Multi-trait analysis of genome-wide association summary statistics using MTAG. Nature Genetics. 2018;50(2):229–37.

15. Wu Y, Cao H, Baranova A, Huang H, Li S, Cai L, et al. Multi-trait analysis for genome-wide association study of five psychiatric disorders. Transl Psychiatry. 2020;10(1):209.

16. Deak JD, Zhou H, Galimberti M, Levey DF, Wendt FR, Sanchez-Roige S, et al. Genomewide association study in individuals of European and African ancestry and multi-trait analysis of opioid use disorder identifies 19 independent genome-wide significant risk loci. Molecular Psychiatry. 2022.

17. Xu H, Toikumo S, Crist RC, Glogowska K, Deak JD, Gelernter J, et al. Multi-trait Analysis of GWAS (MTAG) of Substance Use Traits Identifies Novel Genetic Loci and Phenomic Associations. medRxiv. 2022:2022.07.06.22277340.

18. Foley CN, Staley JR, Breen PG, Sun BB, Kirk PDW, Burgess S, et al. A fast and efficient colocalization algorithm for identifying shared genetic risk factors across multiple traits. Nature Communications. 2021;12(1):764.

19. Abraham G, Qiu Y, Inouye M. FlashPCA2: principal component analysis of Biobank-scale genotype datasets. Bioinformatics. 2017;33(17):2776–8.

20. Fang H, Hui Q, Lynch J, Honerlaw J, Assimes TL, Huang J, et al. Harmonizing Genetic Ancestry and Self-identified Race/Ethnicity in Genome-wide Association Studies. Am J Hum Genet. 2019;105(4):763–72.

21. Justice AC, Smith RV, Tate JP, McGinnis K, Xu K, Becker WC, et al. AUDIT-C and ICD codes as phenotypes for harmful alcohol use: association with ADH1B polymorphisms in two US populations. Addiction. 2018;113(12):2214–24.

22. Gaziano JM, Concato J, Brophy M, Fiore L, Pyarajan S, Breeling J, et al. Million Veteran Program: A mega-biobank to study genetic influences on health and disease. J Clin Epidemiol. 2016;70:214–23.

23. Auton A, Brooks LD, Durbin RM, Garrison EP, Kang HM, Korbel JO, et al. A global reference for human genetic variation. Nature. 2015;526(7571):68–74.

24. Das S, Forer L, Schönherr S, Sidore C, Locke AE, Kwong A, et al. Next-generation genotype imputation service and methods. Nat Genet. 2016;48(10):1284–7.

25. Purcell S, Neale B, Todd-Brown K, Thomas L, Ferreira MA, Bender D, et al. PLINK: a tool set for whole-genome association and population-based linkage analyses. Am J Hum Genet. 2007;81(3):559–75.

26. Pasman JA, Verweij KJH, Gerring Z, Stringer S, Sanchez-Roige S, Treur JL, et al. GWAS of lifetime cannabis use reveals new risk loci, genetic overlap with psychiatric traits, and a causal influence of schizophrenia. Nat Neurosci. 2018;21(9):1161–70.

27. Biological insights from 108 schizophrenia-associated genetic loci. Nature. 2014;511(7510):421–7.

28. Wang K, Li M, Hakonarson H. ANNOVAR: functional annotation of genetic variants from high-throughput sequencing data. Nucleic Acids Res. 2010;38(16):e164.

29. Berisa T, Pickrell JK. Approximately independent linkage disequilibrium blocks in human populations. Bioinformatics. 2016;32(2):283–5.

30. Zhang Y, Lu Q, Ye Y, Huang K, Liu W, Wu Y, et al. SUPERGNOVA: local genetic correlation analysis reveals heterogeneous etiologic sharing of complex traits. Genome Biol. 2021;22(1):262.

31. Abecasis GR, Auton A, Brooks LD, DePristo MA, Durbin RM, Handsaker RE, et al. An integrated map of genetic variation from 1,092 human genomes. Nature. 2012;491(7422):56–65.

32. Pruim RJ, Welch RP, Sanna S, Teslovich TM, Chines PS, Gliedt TP, et al. LocusZoom: regional visualization of genome-wide association scan results. Bioinformatics. 2010;26(18):2336–7.

33. Bulik-Sullivan BK, Loh PR, Finucane HK, Ripke S, Yang J, Patterson N, et al. LD Score regression distinguishes confounding from polygenicity in genome-wide association studies. Nat Genet. 2015;47(3):291–5.

34. Lu Q, Powles RL, Abdallah S, Ou D, Wang Q, Hu Y, et al. Systematic tissue-specific functional annotation of the human genome highlights immune-related DNA elements for lateonset Alzheimer’s disease. PLoS Genet. 2017;13(7):e1006933.

35. Watanabe K, Taskesen E, van Bochoven A, Posthuma D. Functional mapping and annotation of genetic associations with FUMA. Nature Communications. 2017;8(1):1826.

36. Schmitt AD, Hu M, Jung I, Xu Z, Qiu Y, Tan CL, et al. A Compendium of Chromatin Contact Maps Reveals Spatially Active Regions in the Human Genome. Cell Rep. 2016;17(8):2042–59.

37. Yang C, Li S, Ma JX, Li Y, Zhang A, Sun N, et al. Whole Exome Sequencing Identifies a Novel Predisposing Gene, MAPKAP1, for Familial Mixed Mood Disorder. Front Genet. 2019;10:74.

38. Baurley JW, Edlund CK, Pardamean CI, Conti DV, Krasnow R, Javitz HS, et al. Genome-Wide Association of the Laboratory-Based Nicotine Metabolite Ratio in Three Ancestries. Nicotine Tob Res. 2016;18(9):1837–44.

39. Kember RL, Vickers-Smith R, Zhou H, Xu H, Dao C, Justice AC, et al. Genetic underpinnings of the transition from alcohol consumption to alcohol use disorder: shared and unique genetic architectures in a cross-ancestry sample. medRxiv. 2021:2021.09.08.21263302.

40. Schumann G, Liu C, O’Reilly P, Gao H, Song P, Xu B, et al. KLB is associated with alcohol drinking, and its gene product β-Klotho is necessary for FGF21 regulation of alcohol preference. Proc Natl Acad Sci U S A. 2016;113(50):14372–7.

41. Howard DM, Adams MJ, Clarke TK, Hafferty JD, Gibson J, Shirali M, et al. Genome-wide meta-analysis of depression identifies 102 independent variants and highlights the importance of the prefrontal brain regions. Nat Neurosci. 2019;22(3):343–52.

42. Wootton RE, Richmond RC, Stuijfzand BG, Lawn RB, Sallis HM, Taylor GMJ, et al. Evidence for causal effects of lifetime smoking on risk for depression and schizophrenia: a Mendelian randomisation study. Psychol Med. 2020;50(14):2435–43.

43. Teslovich TM, Musunuru K, Smith AV, Edmondson AC, Stylianou IM, Koseki M, et al. Biological, clinical and population relevance of 95 loci for blood lipids. Nature. 2010;466(7307):707–13.

44. Costas J. The highly pleiotropic gene SLC39A8 as an opportunity to gain insight into the molecular pathogenesis of schizophrenia. American Journal of Medical Genetics Part B: Neuropsychiatric Genetics. 2018;177(2):274–83.

45. de Vries PS, Brown MR, Bentley AR, Sung YJ, Winkler TW, Ntalla I, et al. Multiancestry Genome-Wide Association Study of Lipid Levels Incorporating Gene-Alcohol Interactions. Am J Epidemiol. 2019;188(6):1033–54.

46. Gaddis N, Mathur R, Marks J, Zhou L, Quach B, Waldrop A, et al. Multi-trait genome-wide association study of opioid addiction: <em>OPRM1</em> and Beyond. medRxiv. 2021:2021.09.13.21263503.

47. Kember RL, Vickers-Smith R, Xu H, Toikumo S, Niarchou M, Zhou H, et al. Cross-ancestry meta-analysis of opioid use disorder uncovers novel loci with predominant effects in brain regions associated with addiction. Nature Neuroscience. 2022.

48. Weinberger AH, Pacek LR, Giovenco D, Galea S, Zvolensky MJ, Gbedemah M, et al. Cigarette Use Among Individuals with Alcohol Use Disorders in the United States, 2002 to 2016: Trends Overall and by Race/Ethnicity. Alcohol Clin Exp Res. 2019;43(1):79–90.

49. Gulliver SB, Kamholz BW, Helstrom AW. Smoking cessation and alcohol abstinence: what do the data tell us? Alcohol Res Health. 2006;29(3):208–12.

50. Masaoka H, Ito H, Gallus S, Watanabe M, Yokomizo A, Eto M, et al. Combination of ALDH2 and ADH1B polymorphisms is associated with smoking initiation: A large-scale crosssectional study in a Japanese population. Drug Alcohol Depend. 2017;173:85–91.

51. Fawcett KA, Barroso I. The genetics of obesity: FTO leads the way. Trends Genet. 2010;26(6):266–74.

52. Dao C, Zhou H, Small A, Gordon KS, Li B, Kember RL, et al. The impact of removing former drinkers from genome-wide association studies of AUDIT-C. Addiction. 2021;116(11):3044–54.

