## Supplementary Figures for "Multi-trait genome-wide association analyses leveraging alcohol use disorder findings identify novel loci for smoking behaviors in the Million Veteran Program"

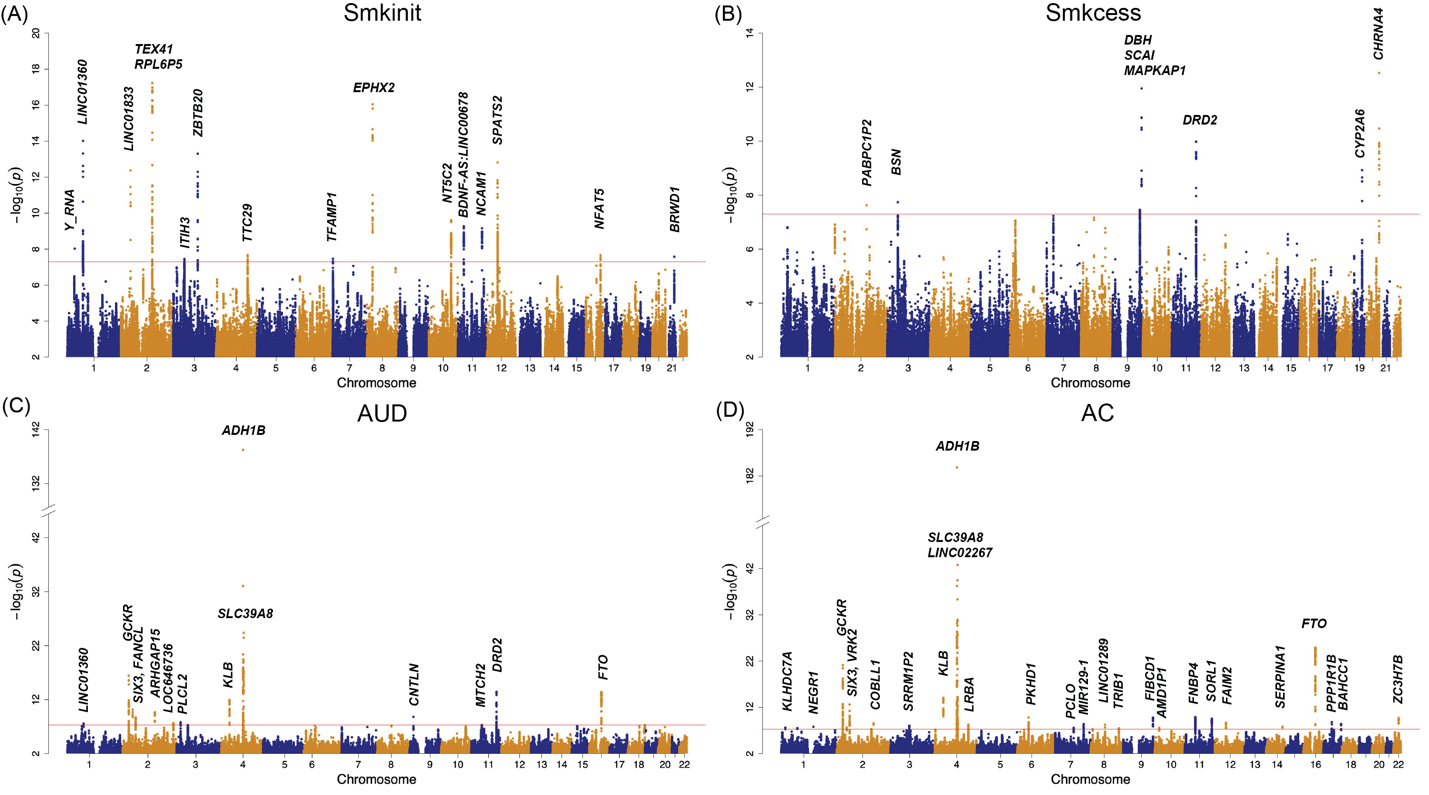

**Supplementary Figure 1: Single-trait genome-wide association study (GWAS) on smoking and alcohol phenotypes.** Manhattan plot of single-trait GWAS for (A) smoking initiation, (B) smoking cessation, (C) AUD, and (D) AC. The nearest genes for genome-wide significant (GWS) loci are shown.

Smkinit: smoking initiation; Smkcess: smoking cessation; AUD: alcohol use disorder; AC: alcohol consumption.

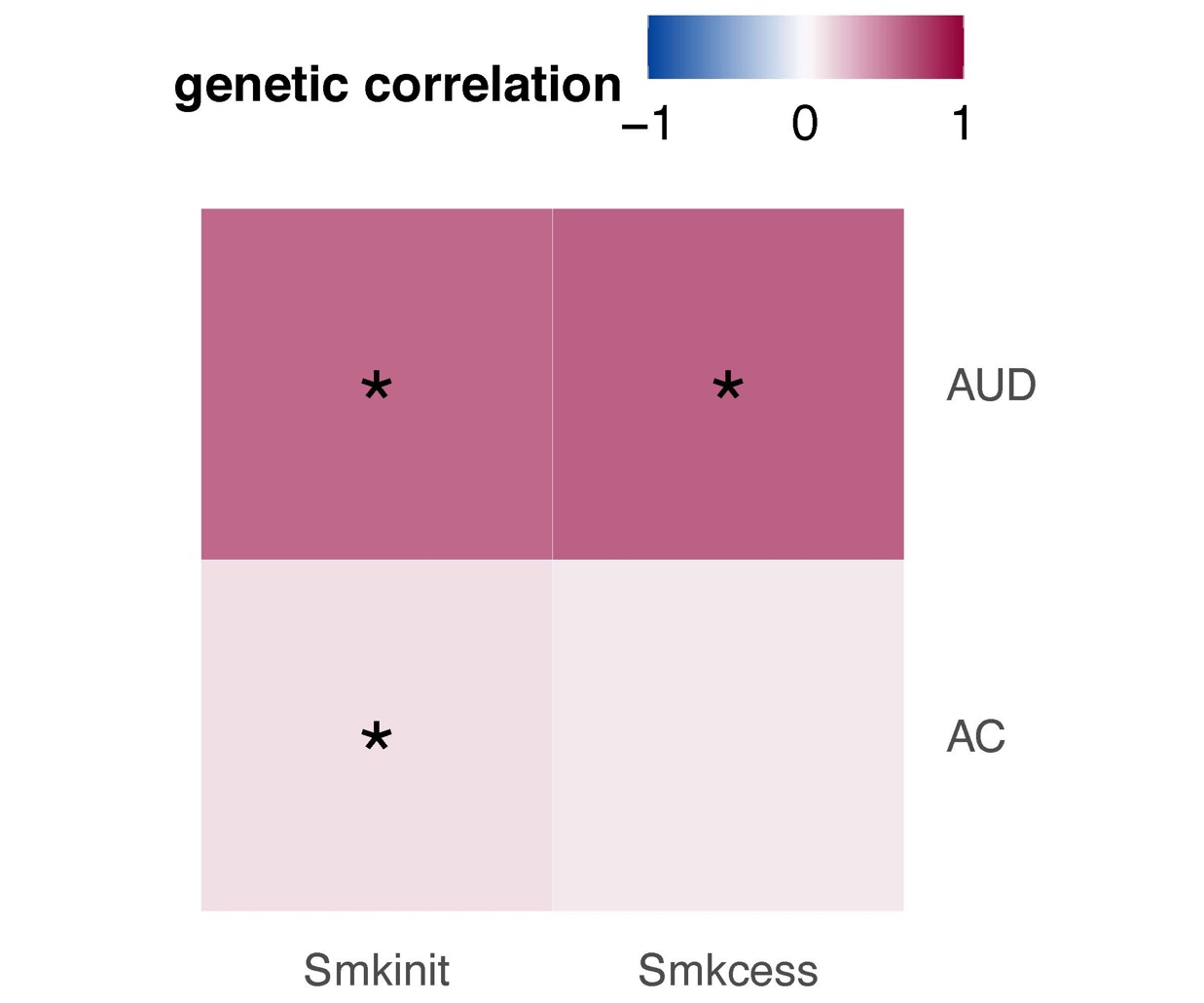

**Supplementary Figure 2: Genetic correlation between smoking and alcohol traits estimated from single-trait GWAS.** Genetic correlations of one smoking trait with one alcohol trait were estimated. The darkness of color indicates the magnitude of the genetic correlation, and the asterisk (*) indicates significance. The genetic correlation was considered significant if p < 0.05/4 = 0.0125 (Bonferroni correction).

Smkinit: smoking initiation; Smkcess: smoking cessation; AUD: alcohol use disorder; AC: alcohol consumption.

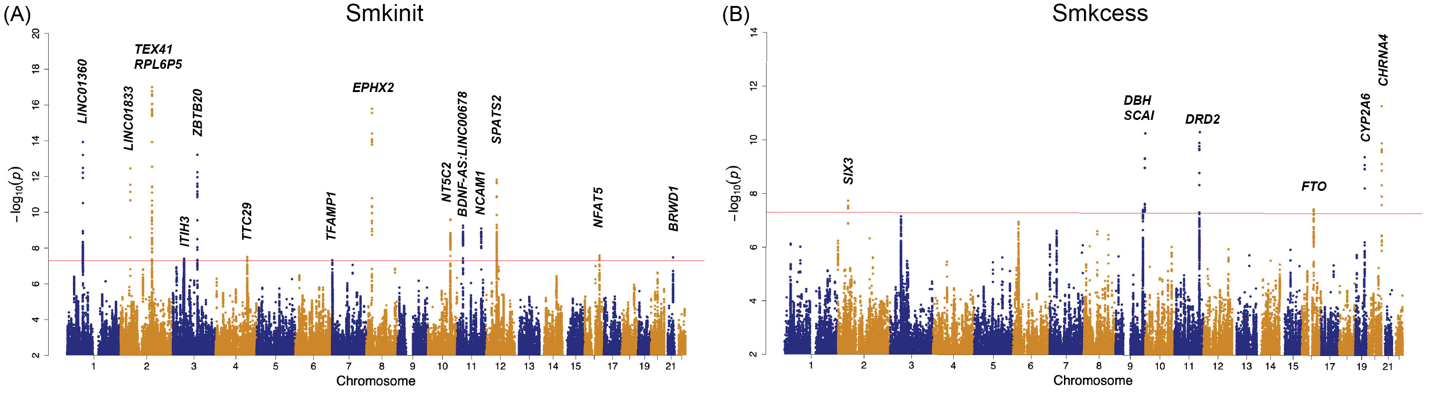

**Supplementary Figure 3: Multi-trait analysis of GWASs (MTAG) on two smoking phenotypes with alcohol consumption (AC).** Manhattan plot of MTAG with AC for (A) smoking initiation and (B) smoking cessation. The nearest genes to GWS loci are shown.

Smkinit: smoking initiation; Smkcess: smoking cessation; AUD: alcohol use disorder; AC: alcohol consumption.

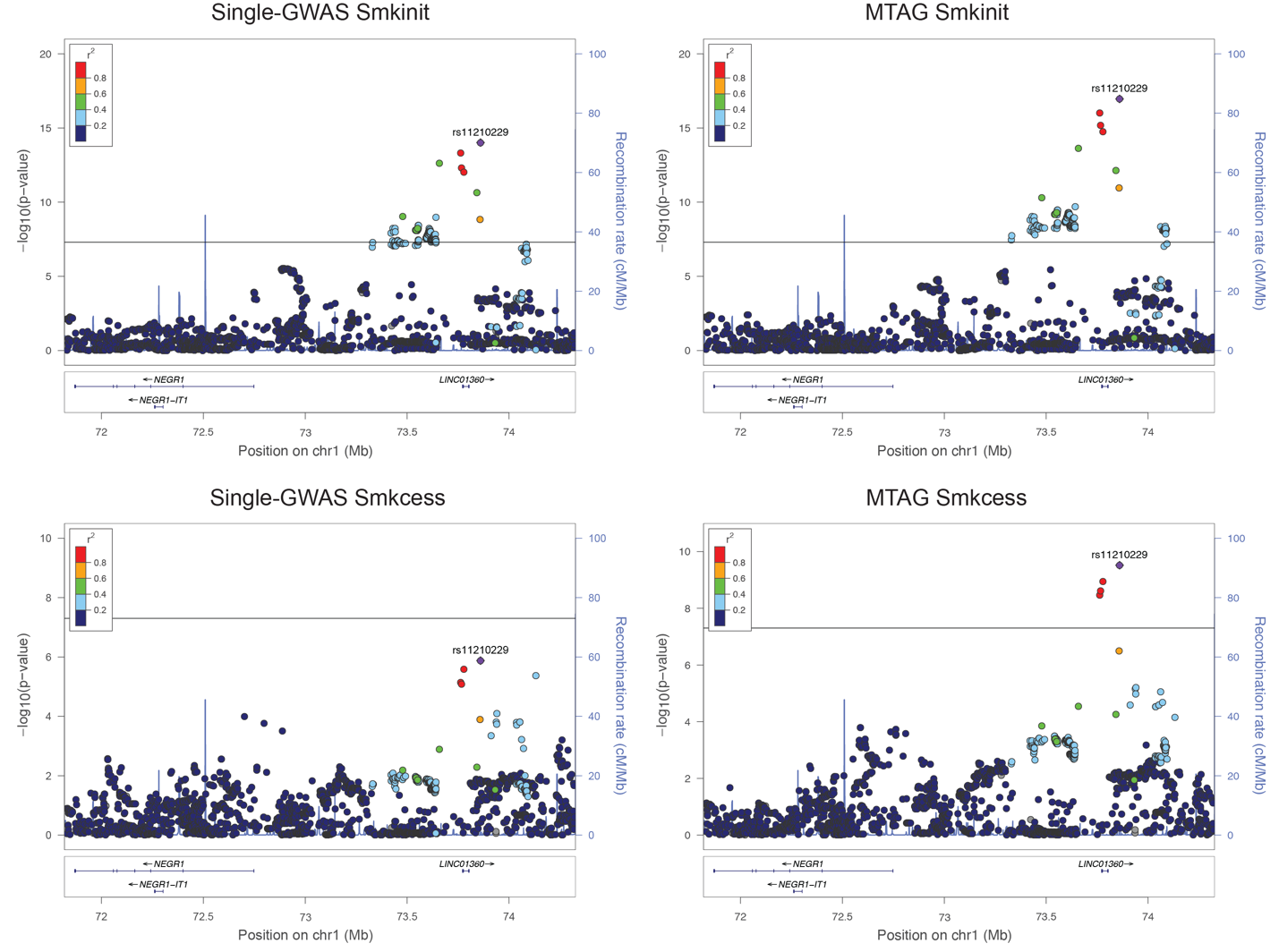

**Supplementary Figure 4: Stacked association plots of** **rs11210229 (*LINC01360*) in the 2 smoking-related GWASs.** The genetic variant rs11210229 mapped on *LINC01360* was identified as a colocalized SNP between AUD and 2 smoking related traits.

Smkinit: smoking initiation; Smkcess: smoking cessation; AUD: alcohol use disorder.

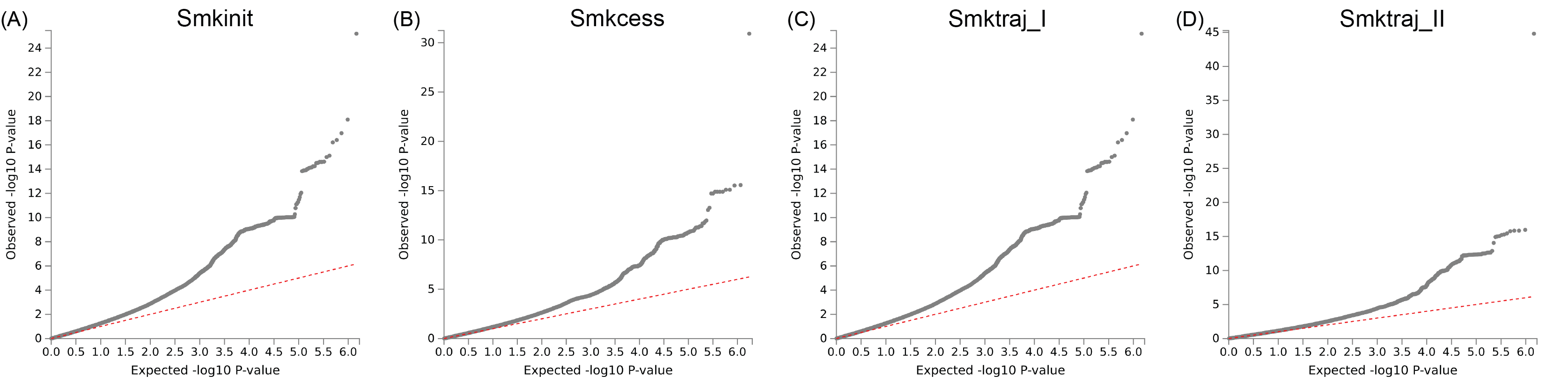

**Supplementary Figure 5: Quantile‒quantile (QQ) plots for the GWAS of 2 smoking related traits after MTAG with AUD.** QQ plot to reflect the increased significance of GWAS results for (A) smoking initiation and (B) smoking cessation.

Smkinit: smoking initiation; Smkcess: smoking cessation; AUD: alcohol use disorder.
